# BMP4 induces a p21-dependent cell state shift in glioblastoma linking mesenchymal transition to senescence

**DOI:** 10.1101/2024.06.20.599819

**Authors:** Mia Niklasson, Erika Dalmo, Anna Segerman, Veronica Rendo, Bengt Westermark

## Abstract

Bone morphogenetic protein 4 (BMP4) has emerged as a potential glioblastoma therapy due to its anti-proliferative effect via SOX2 downregulation and differentiation promotion. However, BMP4 responses vary across and within tumors. Our previous data indicate that BMP4 induces transition to a mesenchymal-like cell state. Mesenchymal transition is associated with therapy-resistance and tumor recurrence, as is senescence in cancer.

In this study, we investigated BMP4’s potential to induce senescence in primary glioblastoma cells, including proneural- and mesenchymal-like clones derived from the same tumor. BMP4 treatment induced senescence-associated genes and phenotypic changes such as cell enlargement, senescence-associated-β-gal expression, lamin B1 downregulation, and elevated p21 levels. The most robust senescence induction was observed in the mesenchymal-like clone, compared to its proneural counterpart. Notably, mesenchymal-like cells displayed high basal levels of p21 and other senescence-associated markers, suggesting a convergence of mesenchymal and senescent traits. p21 knockout abolished BMP4-induced senescence, maintaining proliferation and cell size despite SOX2 downregulation. Additionally, senolytic treatment effectively eliminated senescent cells through apoptosis, thereby favoring survival of cells retaining normal p21 levels.

Our findings demonstrate BMP4’s ability to induce p21-dependent senescence in glioblastoma, particularly in therapy-resistant mesenchymal-like cells. These insights provide potential therapeutic strategies targeting senescence pathways in this challenging disease.

## Introduction

Glioblastoma (GBM) is the most malignant brain cancer in adults. Despite rigorous standard-of-care interventions, including surgical resection and concomitant radio- and chemotherapy, the median survival of patients is merely around 15 months from diagnosis. This unfavorable prognosis is attributed, in part, to the significant intra- and intertumoral heterogeneity.

The extensive intratumoral heterogeneity observed in GBM is partly ascribed to significant cellular plasticity. Malignant cells can transition between various cell states, primarily along a gradient between the neurodevelopmental/proneural (PN) and injury response/mesenchymal (MES) gene programs (1–3). Notably, the MES cell state is closely associated with heightened therapy-resistance (4) and a reactive/inflammatory profile (5, 6). Recurrent tumors often show mesenchymal characteristics (7), which may be due to survival and expansion of therapy-resistant cells and/or that therapy induces a more MES-like phenotype, so called mesenchymal transition (8, 9).

The inflammatory profile of MES-like GBM cells shows similarities with that of senescent cells, including the release of inflammatory cytokines, extracellular matrix components and proteases, altogether forming the senescence-associated secretory phenotype (SASP). Cellular senescence is generally defined as a state of permanent cell cycle arrest that serves as a protective mechanism to prevent the proliferation of transformed cells and stimulate immune clearance via SASP. Senescence is often characterized by senescence-associated β-galactosidase (SA-β-gal) expression, heightened metabolic activity, increased cell size, and downregulated lamin B1 (10–12). Replicative senescence is associated with telomere shortening (13), but senescence can also be triggered by other factors, such as prolonged activation of oncogenes, exposure to irradiation, oxidative stress, and chemotherapeutic agents like doxorubicin and cisplatin (14–16). Abnormal cellular size increase, leading to cytoplasmic dilution and an imbalanced proteome, can also trigger senescence (17–19). Additionally, certain cytokines can induce senescence in an autocrine or paracrine manner (20).

While cellular senescence is widely recognized as anti-tumorigenic, growing evidence supports the notion that senescent cells can paradoxically promote tumor growth. This can occur either through intrinsic reprogramming of senescent cancer cells into a more stem-like state, allowing them to re- enter the cell cycle (21), or through the secretion of factors such as cytokines, which affect the surrounding cells (reviewed in (22)).

Cells can display reversible markers of senescence under certain conditions, such as after exposure to various cytokines. In particular, the effect of transforming growth factor beta (TGFβ) on cellular senescence is well-documented (reviewed in (23)), but there are surprisingly few reports on the influence of other TGFβ family members, such as the bone morphogenetic proteins (BMPs). BMPs play a crucial role in neurogenesis regulation during both developmental stages and in the adult brain (24, 25). In the aging brain, the concentration of BMP4 has been shown to increase, contributing to compromised neurogenesis and cognitive function (26). In GBM, BMP4 treatment has been demonstrated to downregulate stemness markers such as SOX2, inhibit proliferation (27), and induce (a reversible) astrocytic differentiation (28–30). These pro-differentiation and anti-tumorigenic effects of BMP4 led to a clinical phase I trial where approximately 20% of the patients exhibited partial or complete responses (31). This aligns with the considerable variability in BMP4 response among cell cultures from patient tumors (27, 32).

Furthermore, recent findings indicate significant heterogeneity in BMP4 response within the GBM cell culture, with low BMP4 doses prompting transitions into distinct cell states, including mesenchymal, quiescence, and stem-like states (3). While BMP4 has been reported to induce quiescence in GBM cells (33), it remains crucial to investigate whether BMP4 can induce senescence.

In the current study we set out to expand the knowledge of the heterogeneous response of GBM cells to BMP4, focusing on the potential of BMP4 to induce senescence. We investigate how senescence is related to mesenchymal gene expression, and whether PN- and MES-like GBM cells from the same tumor respond differently to BMP4. We find that BMP4-induced senescence depends on SMAD signaling and p21 upregulation. This transition to a senescent state is more prominent in a MES-like clone compared to a PN-like clone from the same tumor. Finally, we show that senolytic treatment specifically depletes the senescent cell population by inducing apoptosis. This study highlights a previously unrecognized effect of BMP4, namely, the induction of a senescence-like phenotype and its association with the mesenchymal cell state in glioblastoma. These findings emphasize the importance of gaining a nuanced understanding of BMP4’s effects on glioblastoma cells when implementing it in therapeutic interventions.

## Results

### The BMP4-SMAD signaling pathway induces a senescence-like phenotype in GBM cell cultures

We first explored the role of BMP4 in inducing senescence in the primary human GBM cell line U3065MG. Our previous work has already showcased this cell line’s heterogeneity and extensive plasticity, which we accomplished using both single-cell RNA sequencing of barcoded cells (3) and the derivation of phenotypically different clonal cultures (4). Treatment with recombinant human BMP4 (10 ng/ml) for up to two weeks resulted in a reduction in cell proliferation rate (Fig. 1A), consistent with our previous findings (27). To explore global transcriptional changes associated with this phenotype, we performed bulk RNA sequencing and Gene Set Enrichment Analysis (GSEA) using the Chemical and Genetic Perturbation module. Interestingly, gene sets connected to cellular senescence and the mesenchymal GBM subtype were found among the top 20 enriched gene sets (out of 3,858 gene sets) in BMP4-treated cells. This shift was coupled to a reduction of proliferation-associated transcripts and a proneural GBM profile (Supplementary Table 1).

**Figure 1.**
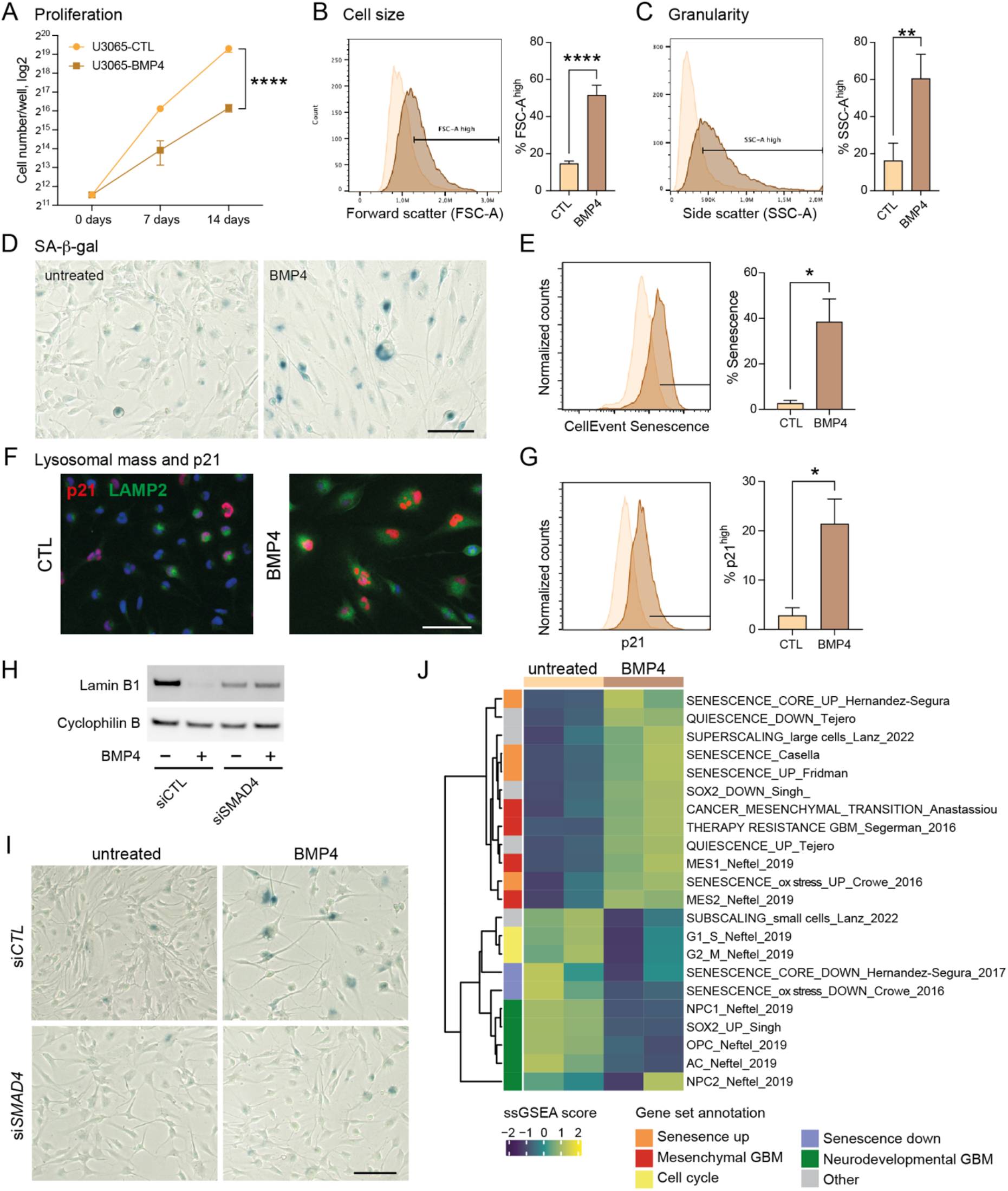
The BMP4-SMAD signaling pathway induces cell size enlargement and a senescence-like phenotype in GBM cell cultures. A. Proliferation rate of U3065MG cells (cell counting on day 0, 7 and 14), untreated or treated with BMP4 (10ng/ml). Unpaired t-test, ****, p<0.0001. B-C. Histograms (left) showing cell size measurement (B) and granularity (C) using flow cytometry forward scatter (FSC-A) and side scatter (SSC-A) area values, respectively, and quantification (right) of the FSC-A and SSC-A high populations of quadruplicate experiments. Unpaired t-test, ****, p<0.0001; **, p=0.0015. D. Photographs of U3065MG cells +/-BMP4 (14 days) stained for SA-β-gal, scale bar 100 µm. E. Histograms of cells stained for SA-β-gal (CellEventSenescence kit) and analyzed by flow cytometry, left, and quantification of the CellEventSenescence-high cell population (gate in histogram) from two experiments, right. *, p≤0.05. F. p21 and lysosomal marker LAMP2 immunofluorescence staining. G. Histograms of cells stained with p21 antibody and analyzed by flow cytometry, left, and quantification of highly p21-expressing cells (two experiments), right. *, p≤0.05. H. Lamin B1 protein levels in control (siCTL) and SMAD4 knockdown (siSMAD4) cells +/-BMP4 (14 days) using western blot analysis. Cyclophilin B was used as loading control. I. SA-β-gal staining of knockdown experiment, scale bar 100 µm. J. Heatmap of single-sample Gene Set Enrichment Analysis (ssGSEA) using senescence- and GBM-related gene sets on transcriptome data from untreated or BMP4-treated (14 days) U3065MG cells, two experiments.

To validate the induction of senescence by BMP4, we conducted a series of senescence-related assays. Unlike quiescent cells, senescent cells typically exhibit abnormal cellular growth, i.e., cell size enlargement, and an increase in granularity, both of which we observed following BMP4 treatment (Fig. 1B-C, respectively). Furthermore, staining for senescence-associated (SA)-β-gal revealed an induction in approximately 40% of the cells (Fig. 1D and 1E). Notably, the extensive cell enlargement was observed within one week’s treatment and preceded the expression of SA-β-gal.

Senescence can be triggered by different signaling pathways, mainly p16^INK4a^-RB and p53-p21^Waf1/Cip1^. In the context of GBMs, it’s noteworthy that approximately 60% of these tumors harbor deletions or alterations in the *CDKN2A* gene (encoding the p16^INK4a^ protein) (cbioportal.org), which also applies for the U3065MG cell line. We investigated the expression of the cyclin-dependent kinase (CDK) inhibitor p21^Waf1/Cip1^ (p21) and indeed observed elevated levels of p21 in BMP4-treated cells in comparison to untreated cells (Fig. 1F and 1G). Moreover, BMP4 treatment induced an increase in lysosomal mass, as evidenced by LAMP2 staining (Fig. 1F), and downregulated the nuclear membrane protein lamin B1 (Fig. 1H), representing additional features of senescent cells (10, 35).

Bound to its receptor, BMP4 can signal via both canonical (involving SMAD proteins) and non-canonical signaling pathways. In U3065MG cells, a one-hour BMP4 stimulation elicited robust phosphorylation of SMAD1/5/9 (Supplementary Fig. 1A). To ascertain the pivotal role of the canonical pathway in the induction of a senescence-like phenotype, we conducted siRNA-mediated knockdown of *SMAD4* during two weeks. In siRNA control cells, BMP4 treatment prompted the upregulation of the senescence-related transcription factor *CEBPB* and the SASP-related *IGFBP7* (Supplementary Fig. 1B) (36, 37), which was not observed in *SMAD4* knockdown cells. Furthermore, *SMAD4* knockdown cells did not exhibit an increase in cell size or granularity in response to BMP4 (Supplementary Fig. 1C-D). Neither did they downregulate lamin B1 (Fig. 1H) or induce SA-β-gal expression (Fig. 1I). Collectively, these results demonstrate the vital role of the canonical SMAD pathway in the induction of the senescence-like phenotype.

We next performed single-sample GSEA (ssGSEA) analyses of transcriptome data from both the U3065MG and the knockdown experiments +/-BMP4, using selected gene sets related to GBM, cell growth, cell cycling, quiescence, and senescence (2, 4, 5, 17, 38–41), as well as MSigDB Hallmarks (Fig. 1J, Supplementary Fig. 1E, and Supplementary Fig. 2). As shown in Figure 1J, BMP4 treatment resulted in enrichment of several gene sets related to senescence induced by various factors (39, 42–44), along with gene signatures related to the mesenchymal GBM subtype, primarily MES1 (2), mesenchymal transition, and multitherapy-resistance (4). In cells with *SMAD4* knockdown, no induction of these signatures was observed with BMP4 treatment (Supplementary Fig. 1E). Since cell enlargement is prevalent—and perhaps even causative—in senescent cells (17, 45), we also used gene signatures connected to cell size, including genes coding for so-called sub-scaling (highly expressed in small cells) and super-scaling (highly expressed in large cells) proteins (17). The dramatic changes of these signatures in BMP4-treated cells clearly support the observed cell size enlargement. We also noted that BMP4 upregulates inflammation-/injury response-related Hallmark gene signatures, such as interferon response, TNFα signaling via NFκB, IL6-JAK-STAT3 signaling, and inflammatory response (Supplementary Fig. 2).

In contrast, BMP4 treatment downregulated signatures related to cell cycling and gene sets annotated as neurodevelopmental GBM, such as activity of the stemness transcription factor SOX2. We also observed downregulation of neurodevelopmental/PN-associated GBM cell state signatures, such as neuronal progenitor cells (NPC1 and NPC2), oligodendrocyte precursor cells (OPC), and notably also astrocyte-like cells (AC) (2, 41) (Fig. 1J). In conclusion, activation of the BMP4-SMAD signaling pathway induces mesenchymal transition and a senescence-like phenotype in a subpopulation of glioblastoma cells.

### Mesenchymal cells exhibit a more pronounced senescence phenotype than proneural cells upon BMP4 treatment

We next asked whether it is a specific cellular subtype or cell state that responds to BMP4 by inducing senescence. To delve deeper into the heterogeneity behind BMP4-mediated induction of senescence, we made use of our previously generated GBM clone libraries. These clone libraries encompass 114 clones derived from five distinct treatment naïve patient tumors, including U3065MG. Clones within tumors exhibit a gradient from therapy-sensitive to therapy-resistant, which is closely connected to a PN-to-MES gradient (4). By re-analysing our previously published transcriptional data originating from the GBM clone libraries, we found a strong correlation between their expression of the senescence signature genes (42) and the expression of a mesenchymal GBM signature (46)(Fig. 2A). This correlation was further supported by analysis of GBM tissue (TCGA) and cell line (HGCC, hgcc.se (34)) data using GSEA (Supplementary Fig 2A). Altogether, these data indicate that the expression profile of tumor cells with a mesenchymal phenotype is in “closer proximity” to senescence than that of cells with a proneural profile.

**Figure 2.**
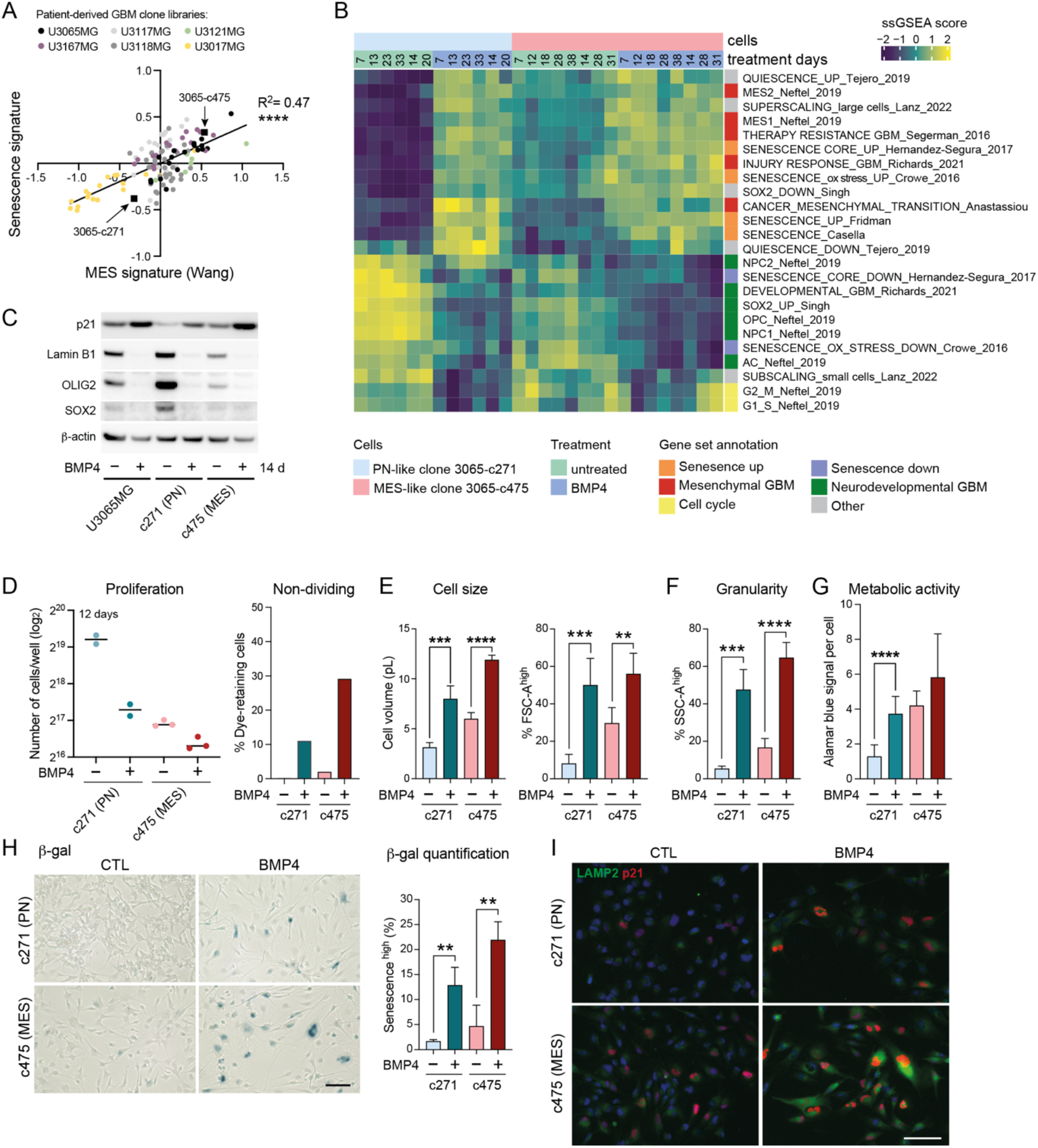
Mesenchymal-like GBM cells display elevated senescence-associated features compared to proneural-like GBM cells. A. Correlation plot of senescence (Fridman, 72 up-regulated genes) and therapy-resistance gene signature scores (Segerman et al) in 114 GIC clones from five patient tumors (U3167MG, U3117MG/U3118MG, U3121MG, U3065MG, and U3017MG). Signature score was calculated by taking the mean of all individual gene z-scores. ****, p=<0.0001. B. Heatmap showing ssGSEA data using gene sets coupled to senescence (39, 42–44), quiescence (40), GBM subtypes (2), GBM multitherapy-resistance (4) and cell size-dependent sub-scaling proteins (17). C. Western blot analysis of the parental cell line U3065MG and its clonal derivatives 3065-c271 (PN-like) and 3065- c475 (MES-like) cells treated +/-BMP4 for 14 days. D. Cell counting of 3065-c271 and 3065-c475 cells treated +/-BMP4 for 12 days (left), and flow cytometry measurement of CellTrace Violet dye- retaining (non-dividing/cell cycle arrested) cells after dye-incubation for the last 5 days (day 8-13, right). E. Cell size measurement using flow cytometry forward scatter (FSC-A) measurements and quantification of the FSC-A high cell population. ** = p≤0.01; *** = p≤0.001. F. Cellular granularity measurement using flow cytometry side scatter (SSC-A) measurements and quantification of the SSC-A high cell population. *** = p≤0.001; **** = p≤0.0001. G. Metabolic activity per cell measured by alamar blue assay in combination with cell counting. H. SA-β-gal staining of cells treated for two weeks +/- BMP4, scale bar 100 µm, (left); and quantification of the CellEventSenescence-high cell population from three experiments (right). ** = p≤0.01. I. p21 and LAMP2 immunofluorescence stainings, scale bar 50 µm.

This finding prompted us to investigate whether PN-like cells exhibit distinct responses to BMP4 treatment compared to MES-like cells. We selected two distinct clones from the parental cell line U3065MG (indicated in Fig. 2A): the PN-like clone 3065-c271 and the MES-like clone 3065-c475 (4, 5). These clones exhibit differential expression of the NPC-related marker CD24 and the MES/inflammatory-related marker CD44 (Supplementary Fig. 3B)(2). The clones were subjected to BMP4 treatment over a period of up to 5.5 weeks, during which RNA was periodically collected for bulk sequencing. An ssGSEA analysis showed that BMP4 in both clones induced deregulation of the same gene sets as in the parental cell line U3065MG (Fig. 1J and 2B), such as gene sets related to senescence, super-scaling, invasiveness, therapy-resistance, and GBM subtypes (Fig. 2B). Moreover, we observed enrichment of genes related to increased cellular metabolism, such as the reactive oxygen species (ROS) pathway, and injury response/inflammation upon BMP4 treatment (6) (Fig. 2B and Supplementary Fig. 3B). An obvious observation was that most of these gene sets had higher baseline levels in the untreated MES-like clone compared to the untreated PN-like clone (Fig. 2B and Supplementary Fig. 3B), indicating a connection between mesenchymal-, senescence- and stress response-related gene expression.

In contrast, BMP4 treatment led to decreased expression of gene sets connected to cell cycling (Fig. 2B and Supplementary Fig. 3C), which was supported by upregulation of cyclin-dependent kinase (CDK) inhibitor p21 protein levels (Fig. 2C and I), indicating cell cycle arrest. This was further confirmed by cell counting experiments, where BMP4 inhibited cell proliferation in both clonal cultures—more so in the fast-proliferating PN-like 3065-c271 than in the MES-like 3065-c475— relative to their untreated controls (Fig. 2D, left). Additionally, a cell tracing dye administered to the clones on treatment day 8 (+/-BMP4) revealed a subpopulation of cells that had not undergone division five days later, supporting the observed inhibition of cell proliferation by BMP4 (Fig. 2D, right). BMP4 has previously been reported to induce cell cycle exit through quiescence (33). Although we indeed observe enrichment of a quiescence-related gene set upon BMP4 treatment, a concomitant induction of genes downregulated in quiescence was observed (40) (Fig. 1J, 2B).

We have previously shown that BMP4-mediated inhibitory effects on proliferation are partially explained by downregulation of the stemness transcription factor SOX2 (27). Here, we observed that BMP4 treatment downregulated SOX2 both at the transcriptome and protein level, and in particular for the SOX2 high-expressing PN-like 3065-c271 (Fig. 2B-C). OLIG2 showed a similar pattern as SOX2, which connected to the decreased expression of NPC1- and OPC-related transcripts (Fig. 2B- C). While BMP4-mediated suppression of the oligodendrocytic lineage has been reported to induce astrocytic differentiation (47), we observed downregulation of the astrocyte-like (AC) gene signature (2) in both clones upon BMP4 treatment, consistent with observations in the parental cell line (Fig. 1J). We therefore investigated the astrocyte marker GFAP at the protein level. While GFAP upregulation could be observed in both the parental cell line and in the PN-like clone, the MES-like 3065-c475 robustly downregulated GFAP following BMP4 treatment (Supplementary Fig. 3D-E). Interestingly, GFAP downregulation has been observed during both replicative and oxidative stress-induced senescence in astrocytes (39).

Although the MES-like clone is larger in cell size than the PN-like clone, BMP4 caused a cell size enlargement of both clones, as shown by enrichment of superscaling genes and increased cell diameters measured in both a cell counter and by flow cytometry (Fig. 2B and E), as well as increased cellular granularity (Fig. 2F). Senescent cells often show an increased metabolic activity compared to both actively dividing cells and quiescent cells, partly due to their high production and secretion of SASP factors. To measure metabolic activity in the cells, we used the alamar blue assay, which measures reduction of resazurin to resorufin in viable cells with an active metabolism, together with cell counting and could observe an increased metabolic activity per cell in BMP4-treated cells (Fig. 2G). To further verify senescence induction by BMP4, cells were analyzed for their expression of SA-β-gal (Fig. 2H), lamin B1 (Fig. 2C), LAMP2 (Fig. 2I) and p21 (Fig. 2C and I). Indeed, in both clones SA-β-gal, p21, and LAMP2 expression increased with BMP4, whereas lamin B1 decreased. However, the most pronounced expression of these markers was observed in the MES-like 3065-c475. Notably, untreated 3065-c475 cells exhibited higher basal levels of SA-β-gal, p21 and LAMP2, along with lower lamin B1 levels compared to untreated 3065-c271 cells (Fig. 2H-I).

Altogether, these results indicate that the MES-like clone already at baseline displays elevated senescence-associated features compared with the PN-like clone, which experimentally confirms the transcriptome analysis of the GBM clone libraries, cell lines and tumors (Fig. 2A and Supplementary Fig. 3A). We therefore hypothesize that the higher basal level of the CDK inhibitor p21^Waf1/Cip1^ observed in connection to cell enlargement, invasion, inflammation, therapy-resistance and senescence in the MES-like 3065-c475 may play an important role in the decision making to transit into a senescent cell state.

### Cell enlargement and induction of a senescence-like phenotype by BMP4 is dependent on p21

To investigate the role of p21^Waf1/Cip1^ in mediating the senescence-inducing effect of BMP4, we used the CRISPR-Cas9 system to knockout the p21 gene *CDKN1A* in the parental cell line U3065MG as well as in the MES-like clone 3065-c475. These cells were selected because of their stronger induction of senescence compared to the p21 low-expressing PN-like clone 3065-c271. From both 3065-c475 and U3065MG, complete *CDKN1A*/p21 knockout cultures (p21-KO) could be generated (Fig. 3A and Supplementary Fig. 4A).

**Figure 3.**
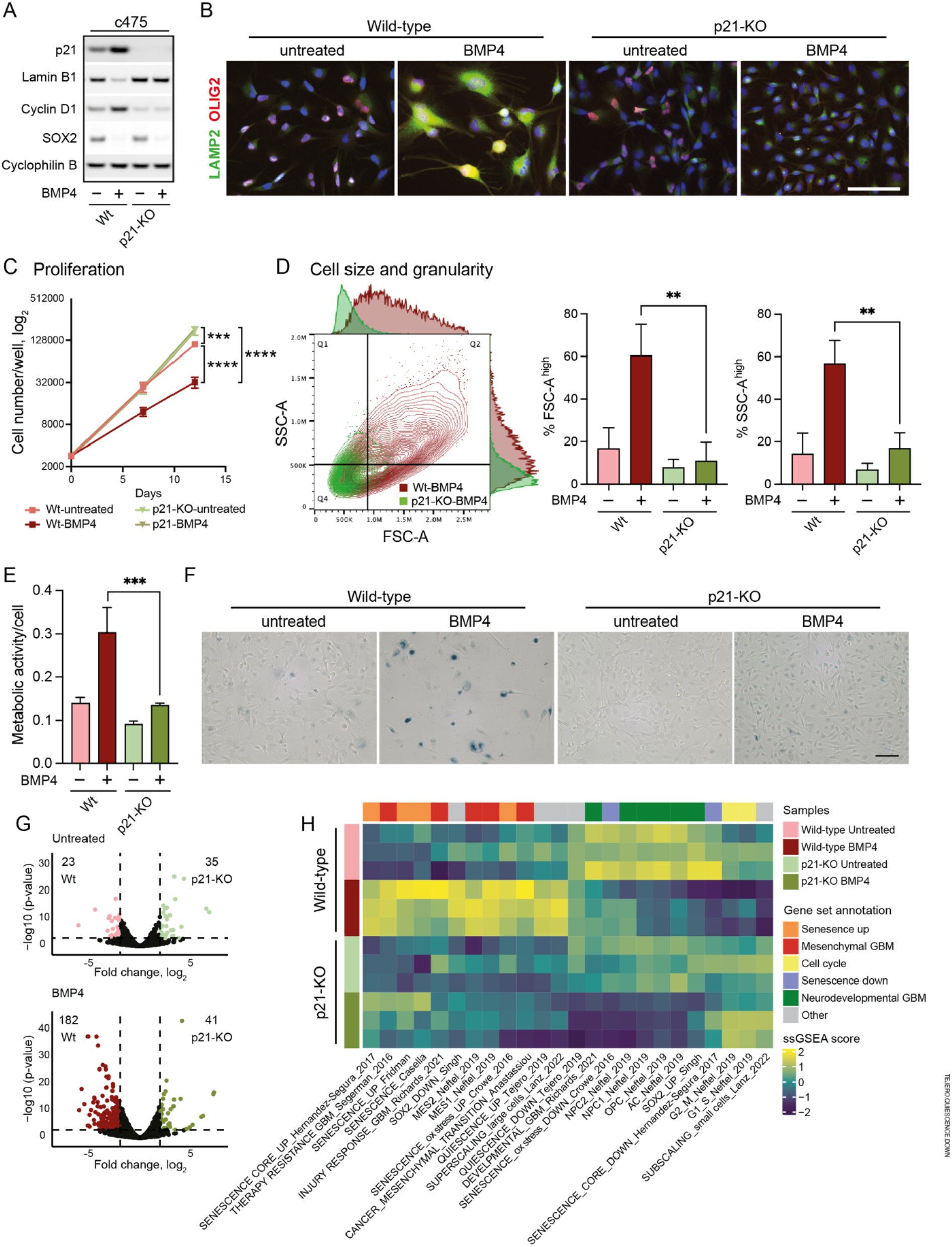
Cell size enlargement and senescence-like induction by BMP4 is dependent on p21. A. Western blot on wild-type and p21 knockout (p21-KO) MES-like clone 3065-c475 cells +/- BMP4 using antibodies against p21, lamin B1, cyclin D1, SOX2, and cyclophilin B. B. Immunocytochemistry of wild-type and p21-KO 3065-c475 cells +/- BMP4 using antibodies against LAMP2 and OLIG2. C. Proliferation of wild-type and p21-KO cells +/- BMP4. Cell counting on day 0, day 7 and day 14. D. Flow cytometry analysis of cell size (cell growth) using forward scatter (FSC-A) measurements and quantification of the FSC-A high cell population; and cell granularity. Side scatter area (SSC-A) high gated population is plotted. The PN-like clone 3065-c271 was used as a reference. E. Metabolic activity (Alamar Blue signal) per individual cell (cell counting) in c475 wild-type and p21-KO cells treated +/- BMP4. F. SA-β-gal staining, scale bar 100 µm. G. Volcano plots of differentially expressed genes (DEGs, 2 ≤ log2 fold difference, p ≤ 0.001) from wild-type and p21 knockout c475 cells, comparing both untreated cells (top, 58 DEGs) and BMP4-treated cells (bottom, 223 DEGs). H. Heatmap of ssGSEA-scores of c475 wild-type and p21-KO cells +/-BMP4. See also Supplementary Figure S4 for data on U3065MG wild-type and p21-KO.

To clarify the functional role of p21 for BMP4-induced senescence, we next performed protein analysis and phenotypic assays. Downregulation of lamin B1 upon BMP4 treatment was essentially abolished in p21-KO cells, whereas the BMP4-induced downregulation of SOX2 occurred also in the knockout. In the absence of p21, cells were no longer driven into G1 cell cycle arrest by BMP4, observed by abolished cyclin D1 upregulation in 3065-c475 p21-KO cells and diminished upregulation in U3065MG cells (Fig. 3A, Supplementary Fig. 4A). Further, BMP4-induced increase in lysosomal mass was lost upon p21 knockout in 3065-c475 (Fig. 3B). Deletion of p21 led to increased cell proliferation rate compared to wild-type cells; this was more pronounced in MES-like 3065-c475 cells than in U3065MG cells. The inhibitory effect of BMP4 on proliferation was attenuated in U3065MG p21-KO cells, whereas in 3065-c475 p21-KO cells it was totally abrogated (Fig. 3C and Supplementary Fig. 4B). In addition, the cellular enlargement induced by BMP4 in wild-type cells was not observed in p21-KO cells, and these cells also exhibited less granularity (Fig. 3D and Supplementary Fig. 4C, D). In 3065-c475 p21-KO cells, the increase of metabolic activity per cell by BMP4 was lost (Fig. 3E). Notably, the BMP4-induced expression of SA-β-gal in wild-type cells was essentially abolished in absence of p21 (Fig. 3F, Supplementary Fig. 4E). In summary, these results show that p21 has a key role in mediating the senescence-like phenotype induced by BMP4 in GBM cells, particularly in more mesenchymal, therapy-resistant cells.

To understand the impact of p21 deletion on transcription, both p21-KO and wild-type cells were treated with BMP4 for two weeks, followed by bulk RNA sequencing and differential expression analysis. Overall, on the transcriptional level, the difference between untreated wild-type and p21-KO cell cultures was moderate (58 differentially expressed genes (DEGs) in 3065-c475, 2 ≤ log2 fold difference, p ≤ 0.001). Upon BMP4 treatment, the number of DEGs between wild-type and p21-KO cells increased substantially to 223 DEGs in 3065-c475 (Fig. 3G). Thus, p21 appears to have a minor role regulating the transcriptional profile of untreated cells, but in BMP4-treated cells the impact of p21 expression is more prominent.

Next, we employed ssGSEA to elucidate the biological differences between BMP4-treated p21-KO cells and BMP4-treated wild-type cells, using the previously selected gene signatures (Fig. 2B). The enrichment of senescence- and MES-GBM-related signatures induced by BMP4 was markedly diminished in p21-KO cells, and there was no induction of the super-scaling gene set (Fig. 3G, Supplementary Fig. 4F). Moreover, the cell cycle- and the sub-scaling-related gene signatures were not affected by BMP4 upon deletion of p21. Interestingly, BMP4 treatment still reduced neurodevelopmental GBM gene signatures irrespective of p21 status, corroborating the observed downregulation of SOX2 and OLIG2 (Fig. 3A-B, Supplementary Fig. 4A). These findings underscore the interconnection among senescence, mesenchymal GBM, and cell size augmentation as responses to BMP4, all mediated via p21 induction. Notably, we found that the induction of the senescence-like phenotype is independent of the reduction of the neurodevelopmental GBM axis, which appears to be linked to SOX2 downregulation.

### Senolytic treatment eradicates BMP4-induced senescent cells and restores mitogenic signaling

To explore whether any preferential therapeutic vulnerabilities exist in high p21-expressing cells, we next performed a systematic correlation analysis between 1,442 cancer cell lines from the Dependency Map dataset [https://depmap.org/portal/] with available RNA expression (log2(TPM+1); 23Q4 public release) and drug sensitivity (area under the curve (AUC); PRISM secondary screen) profiles. This unbiased analysis revealed a significant anticorrelation between *CDKN1A*/p21 baseline expression levels and response to several senolytic agents, suggesting that high p21 expression levels confer sensitivity to treatment (Fig. 4A). Among these drugs, navitoclax (ABT-263)—an inhibitor of anti-apoptotic Bcl-2/-XL/-w proteins—showed the most pronounced effect size (R = −0.13; p=0.027) (Fig. 4A-B). The same trend was observed when focusing on navitoclax response in glioma cell lines only (Fig. 4B). To investigate whether BMP4-treated cells, expressing high p21 levels, are more vulnerable to navitoclax than untreated cells, MES-like 3065-c475 cells (pre-treated +/- BMP4) were exposed to drug treatment for 48 hours. Western blot analysis showed that BMP4 and navitoclax combination treatment leads to decreased p21 levels, comparable to those in untreated controls (Fig. 4C), suggesting an enrichment of the—in terms of senescence—BMP4-non-responding cell population. In line with this, transcriptomic analysis of these cells revealed decreased expression of senescence-related genes (42, 44) compared to BMP4 treatment alone (Supplementary Fig. 5). To validate this, navitoclax-treated 3065-c475 cells +/-BMP4 were fixed and stained for SA-β-gal, resulting in a nearly complete eradication of the SA-β-gal positive cell population (Fig. 4D) and hence an enrichment of the BMP4-non-responding cell population. These cells showed upregulation of cycle-related genes (Supplementary Fig. 5), indicating increased proliferative potential in the non-senescent cell population by navitoclax. Since the Ras-MAPK signaling pathway is recognized as the primary mitogenic pathway in glioblastoma stem-like cultures, immunoblot analysis of phosphorylated ERK1/2 was conducted. Indeed, activation of this pathway was more pronounced in navitoclax- exposed cells, albeit less so in BMP4-treated cells compared to untreated controls (Fig. 4C).

**Figure 4.**
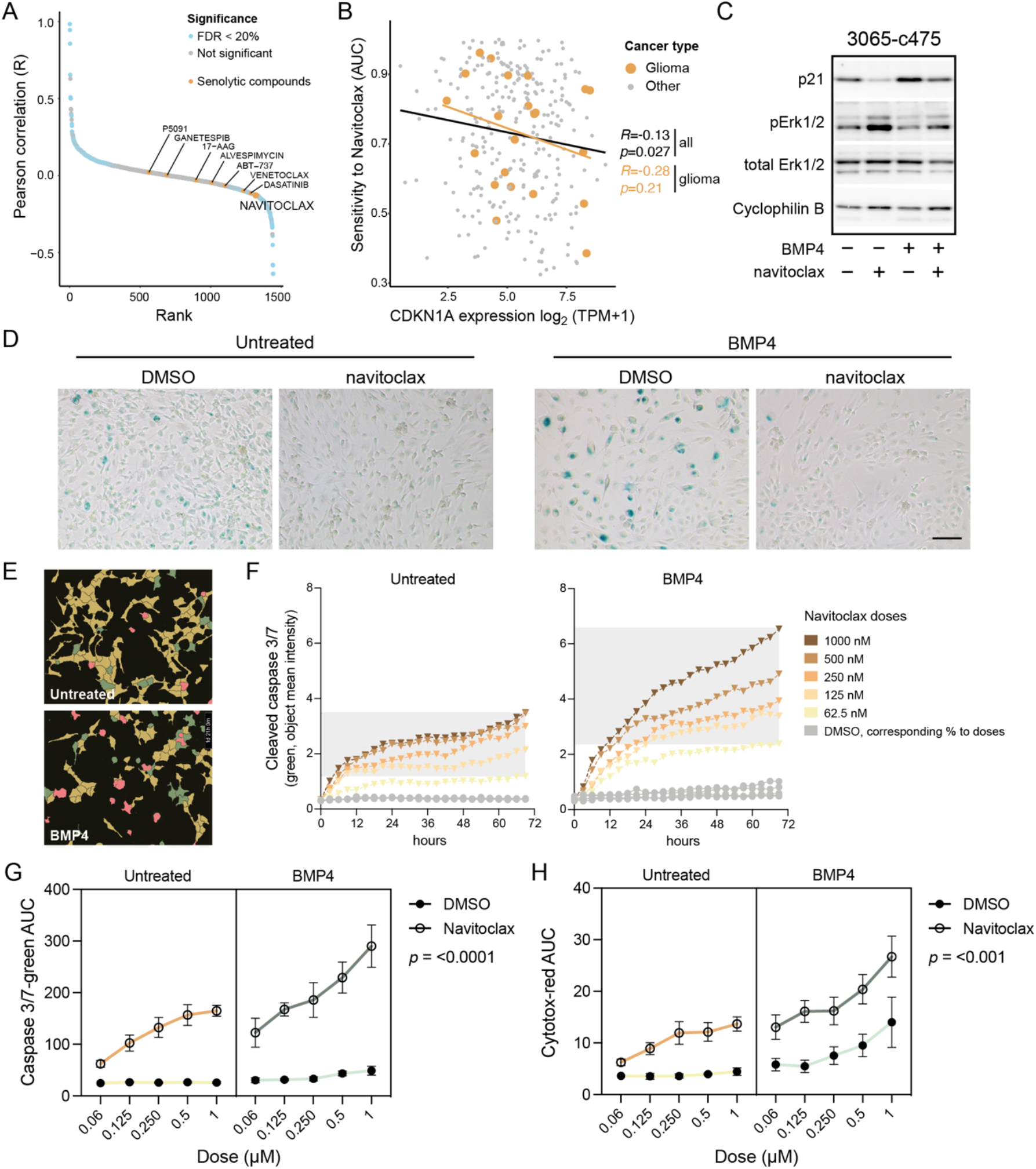
Senolytic treatment eradicates the senescent-like cells via apoptosis. A. Cancer Dependency Map (DepMap) analysis of AUC levels of drugs and compounds and CDKN1A gene expression. Senolytic drugs are indicated in yellow. B. Correlation of CDKN1A expression and navitoclax sensitivity in cancer cell lines. Gliomas are indicated in yellow. C-D. MES-like 3065-c475 cells treated +/-BMP4 for 13 days with navitoclax (0.5-1µM) addition on day 11, analyzed by C) western blot analysis using antibodies against p21, phosphorylated and total ERK1/2, and cyclophilin B1; and D) SA-β-gal staining, scale bar 100 µm. E-H. Navitoclax treatment (five doses, 62,5-1000 nM range) in untreated or BMP4-treated 3065-c475 cells. Navitoclax was added on experimental day 11 and cells were monitored during 72 hours. DMSO-treated cells were used as control and cells were analyzed for activity of cleaved Caspase 3/7 and cytotoxicity. E. Example photographs of cells (125 nM navitoclax for 45 hours) showing masks for individual cell analysis. Red, double-positive cells; green, cleaved caspase 3/7 only; yellow, negative cells. F. Cleaved Caspase 3/7 apoptosis quantification (object mean intensity) of cells. G-H. Three-way ANOVA analysis of treatment at all doses over the 72 hours treatment period measuring cleaved Caspase 3/7 (G) and cytotoxicity (H).

Since navitoclax inhibits the anti-apoptotic machinery active in senescent cells, we next used a cleaved caspase 3/7 assay to assess the apoptotic potential of navitoclax in BMP4-treated cells. Untreated or BMP4-treated 3065-c475 cells received navitoclax or DMSO control in five doses (ranging from 62,5–1000 nM) on treatment day 11. Cells were monitored every third hour over a 72-hour period for cleaved caspase 3/7 activity, as well as cell cytotoxicity (by measuring plasma membrane integrity), followed by cell-by-cell analysis of fluorescence intensities (Fig. 4E). BMP4-treated cells indeed demonstrated increased susceptibility to navitoclax, evidenced by a continuous increase in apoptosis- related fluorescence intensity over the treatment period (Fig. 4F). Area under the curve (AUC) values for each dose during 72 hours (as shown in Fig. 4F) was calculated for both cleaved caspase 3/7 and cytotoxicity assays, followed by three-way ANOVA analyses. The analyses confirmed a substantial difference in navitoclax response—both in terms of apoptosis and cytotoxicity— between untreated and BMP4-treated cells (Fig. 4G-H). Notably, BMP4-treated cells also showed increased cytotoxicity to high doses of DMSO compared to untreated cells, indicating a general decrease in plasma membrane integrity of these cells. In summary, these data demonstrate the efficacy of navitoclax treatment in overcoming BMP4-induced senescence in glioblastoma cells.

## Discussion

Signaling pathways involved in the development of GBM have been extensively studied. There is a consensus that aberrant activation of protein tyrosine kinase receptor signaling and activation of the PI3K/Akt and Ras/MEK pathways are important driver mechanisms, combined with inactivation of the TP53 and CDKN2A/RB pathways (48). Comparatively little is known about inhibitory receptor signaling pathways in GBM. Among these, BMP signaling events have attracted attention, much because of the potential clinical application (reviewed in (49)). A recent clinical phase I trial on recurrent GBM tumors demonstrated that some patients show partial or complete responses to recombinant human BMP4 treatment (31). This finding is remarkable and suggests that context-dependent effects of BMP4 may underlie the differential clinical responses observed. In the present study, we have extended the knowledge on BMP4-mediated effects on GBM cells. Most importantly, we found that canonical BMP4 signaling via p21 induces a senescence-like phenotype, including SA- β-gal expression, downregulation of lamin B1, increased cell size and granularity, and cell cycle arrest. This effect was most pronounced in a mesenchymal-like GBM clone compared to a proneural-like GBM clone. Furthermore, treatment with the senolytic drug navitoclax, a Bcl-2/-XL/-w anti-apoptosis inhibitor, efficiently eliminated the senescent cell population.

BMP4 is a potent inducer of the cell cycle inhibitor p21, prompting us to study its role in the induction of a senescence-like phenotype. Strikingly, we found that deletion of p21 led to an almost complete loss of cell enlargement and SA-β-gal positive cells in BMP4-treated cultures. Additionally, high levels of *CDKN1A*/p21 gene expression were significantly correlated with sensitivity to the senolytic drug navitoclax across multiple cancer cell lines. Conversely, elimination of the SA-β-gal-positive population by navitoclax enriched for cells expressing normal p21 levels. Elevated levels of p21 itself can induce senescence, as shown in several cell lines, including GBM (50, 51). Interestingly, this induction can be prevented by restricting cellular size growth using mTOR inhibitors (50). Since each cell type has an optimal cell size for proper cell function (reviewed in (52)), abnormal cellular enlargement may itself result in senescence (17, 45). Malignant tumors often exhibit a wide variation in cell size, often denoted as pleomorphism, but the underlying mechanisms are poorly understood.

Our study suggests that p21 levels may be involved in cell size regulation in GBM. The relationship between increased cell size, p21 function, and senescence induction remains to be elucidated.

Inhibition of GBM cell proliferation by BMP4 is partially dependent on SOX2 downregulation (27). In the present study, knocking out p21 had no effect on the BMP4-mediated reduction of SOX2 expression. These findings indicate that the cell enlargement and senescence effects of BMP4 are mediated by p21 and not related to SOX2 downregulation (Figure 5). At which point BMP4 signaling is bifurcated remains an open question.

**Figure 5.**
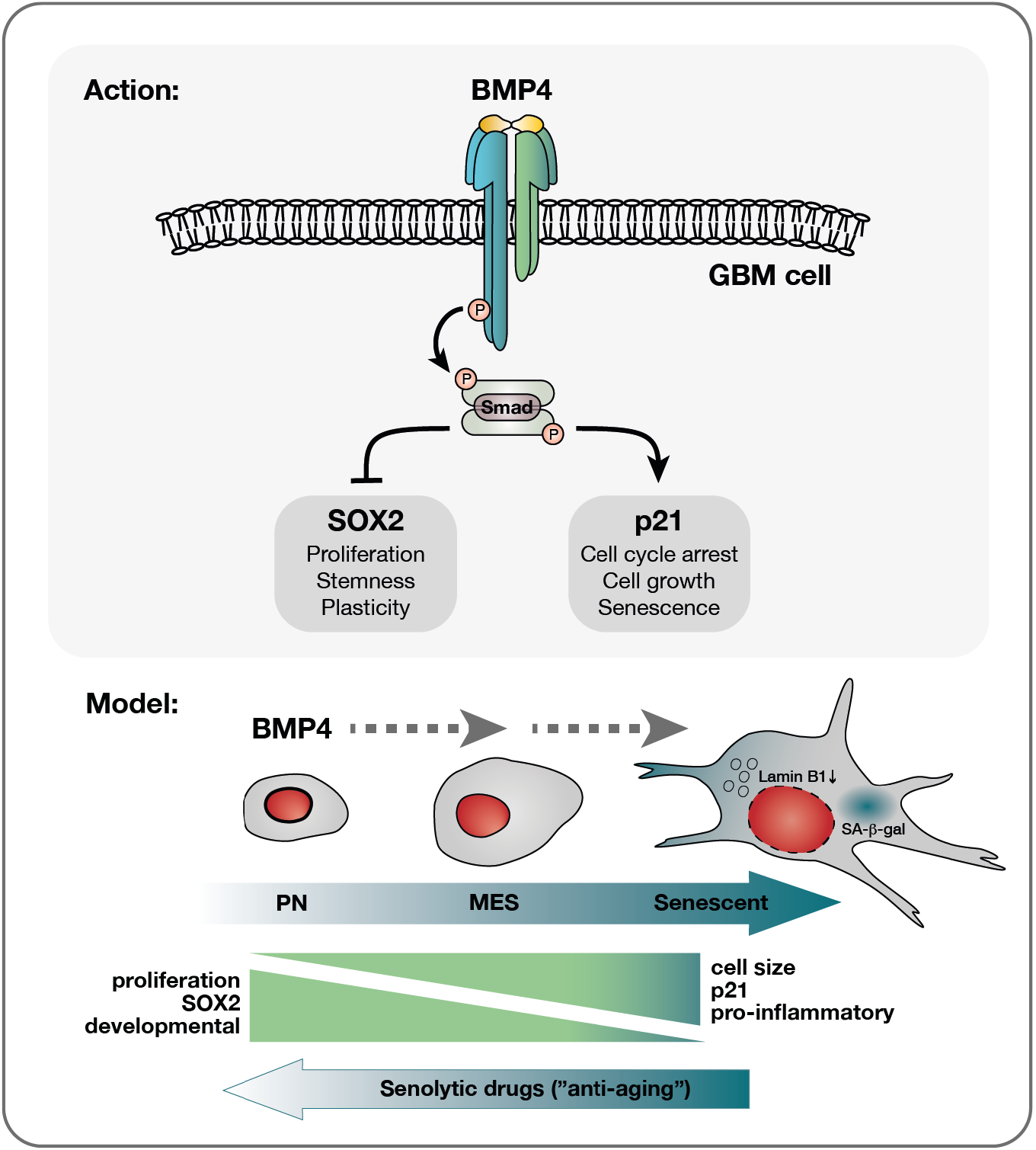
BMP4’s effect on diverse glioblastoma cells reveals a multifaceted response. Upon receptor binding of BMP4, activation of the canonical SMAD complex initiates a bifurcation of the pathway into a SOX2-inhibitory branch and a p21-activating pathway. Consequently, proliferation is inhibited while cellular growth is promoted. This cell cycle arrest yields different outcomes depending on the cellular state: smaller PN-like GBM cells, characterized by lower basal levels of p21 yet higher SOX2 expression, may preferentially opt for differentiation, as evidenced by astrocytic GFAP upregulation (Supplementary Fig. 3D-E). Conversely, larger MES-like GBM cells display a higher propensity to undergo senescence in response to BMP4. This senescence-promoting program is characterized by the upregulation of genes, such as pro-inflammatory genes, which are already expressed at elevated levels in these cells compared to their proneural counterparts. Importantly, given the documented multitherapy-resistant nature of MES-like GBM cells (4), their propensity to enter senescence presents a significant therapeutic opportunity. Here, we demonstrate that BMP4- induced senescent MES-like GBM cells exhibit sensitivity to senolytic treatment, activating apoptotic pathways. Subsequently, the surviving cells post-senolytic treatment display heightened proliferation, thereby rendering them susceptible to conventional therapeutic interventions.

In our previous work, we isolated multitherapy-resistant and -sensitive clones from the early U3065MG cell culture (4), and two of these were used in the current study. Importantly, the therapy-resistant MES-like clone was more vulnerable to BMP4-mediated induction of the senescence-like phenotype than the therapy-sensitive PN-like clone. This vulnerability may be attributed to the MES-like clone’s higher baseline levels of p21, lower lamin B1 expression, and larger cell size—all characteristics associated with senescence. Interestingly, the SOX2 high-expressing PN-like clone responded to BMP4 by inducing astrocytic GFAP, which was not observed in the MES-like clone, and hence we speculate that differentiation versus senescence cell fate, in response to BMP4, may depend on basal levels of SOX2 and p21, respectively.

Resistance to treatment and tumor progression have frequently been associated with mesenchymal transition across various cancers, with recurrent GBM often exhibiting MES characteristics (53). Additionally, the PN-to-MES gradient of malignant cells is molecularly closely related to an increase in an astrocyte reactivity cell state (5), which is part of the CNS injury response. Interestingly, the reactive state includes both cell size enlargement and the induction of inflammatory factors, many of which are part of the SASP. Notably, the MES-like clone used in this study was previously shown to express higher levels of SASP-related proteins, such as CCL2, IL7, IGFBPs, and TIMP-1, compared to the PN-like clone (5). Since senescent cells, like reactive cells, both increase in size and express inflammatory factors, this suggests that GBM cells may exist on a spectrum from a low to high propensity to enter senescence, coupled to the PN–MES gradient (Figure 5). This hypothesis is supported by our pathway analyses of GBM clonal cultures, cell lines, and tissue, which indicate an association between senescence and a MES-like/injury response/therapy-resistant phenotype.

There is a growing understanding of the role of senescent cells in tumor growth (reviewed in (22)). While senescence on one hand plays pivotal roles in impeding the progression of premalignant lesions (16), it may also have adverse effects on the tumor. Recent studies have demonstrated the presence of senescent cells in human GBMs and that a high senescence gene signature score correlates with shorter patient survival (54). Moreover, radiation-induced senescent cells have been shown to promote GBM tumor growth in murine models via the release of SASP factors (55). Thus, the persistent presence of senescence-like cells in glioblastoma bears significant clinical implications, given that the cytokine release from such cells can promote tumor growth (reviewed in (56)). Remarkably, removal of the senescent cells, marked by p16Ink4a expression, led to improved survival in a MES-GBM mouse model (54). Importantly, given the documented multitherapy-resistant nature of mesenchymal-like GBM cells, their propensity to enter senescence that we show in this work, presents a significant therapeutic opportunity. Senolytic therapy therefore holds promise as an adjuvant therapy for GBM patients.

Senescent cells may be generated as a response to intrinsic or extrinsic DNA damaging factors. Our study shows that cells with a senescent phenotype may occur as a response to cytokine signaling, and that this effect may be related to both basal and induced p21 levels. The finding that the senolytic drug navitoclax effectively targets the senescent cell population via apoptosis shows that the survival of these cells depends on anti-apoptotic mechanisms. Further studies on the role of cellular senescence, its connection to mesenchymal transition and therapy resistance, and the effect of senolytic drugs in glioblastoma are highly warranted, both in treatment-naïve tumors and in tumor recurrences.

Importantly, we believe these results have implications beyond GBM and are applicable to other cancers as well.

This study underscores BMP4’s ability to induce senescence in glioblastoma, primarily targeting the most mesenchymal-like cells expressing high p21 levels. These cells demonstrate sensitivity to senolytic treatment, leading to an enrichment of a more proliferative cell population, which could potentially be targeted by conventional therapies.

## Methods

A complete list of Methods can be found in the Supplementary Materials and Methods section.

### Cell culture and treatments

Cells used in this study were the human GBM cell line U3065MG, obtained from the GBM cell line biobank HGCC (hgcc.se, (34)), and clonal cultures from early passage U3065MG (3065-c271 and 3065-c475) (4). Briefly, cells were grown adherently on laminin (Sigma-Aldrich)-coated Primaria cell culture dishes (BD Biosciences) in serum-free neural stem cell media (Neurobasal and DMEM/F12 media (1:1) supplemented with N2, B27 w/o retinoic acid (Thermo Fisher Scientific), EGF (10 ng/ml), and bFGF (10 ng/ml) (Peprotech)). Recombinant human BMP4 (Thermo Fisher Scientific) at 10 ng/ml was added every third day in the presence of EGF and bFGF in the media. During media change, every three to four days, at least 50% of conditioned media was retained. The senolytic drug navitoclax (ABT-263, Selleck Chemicals) was used in 62,5–1000 nM concentrations. See additional information on cell culture and treatments in Supplementary Materials and Methods.

### Beta-galactosidase stainings

Detection of SA-β-gal activity on adherent cells was done using the Senescence β-Galactosidase Staining Kit (#9860, Cell signaling), according to manufacturer’s instructions. For flow cytometry detection of β-gal the CellEvent Senescence Green Flow Cytometry Assay Kit (C10840, Invitrogen) was used according to the recommended protocol, using a 1:1000 probe dilution.

### CRISPR/Cas9 knockout

For knockout of the p21 gene, the p21 Waf1/Cip1 CRISPR/Cas9 KO plasmids (sc-400013, pool of three plasmids with individual 20 nt guide RNA, and a GFP gene) and the p21-specific homology- directed repair (HDR) plasmid containing puromycin resistance gene and RFP gene (sc-400013-HDR) were transfected into cells according to the manufacturer’s recommendations (Santa Cruz). Control CRISPR/Cas9 plasmid (sc-41822, non-targeting 20 nt scramble guide RNA) and Transfection Reagent (TFR) only were used as controls in MES-like clone 3065-c475 and U3065MG, respectively. After 4 days, puromycin (0.5 µg/ml) was added to the cells to select for successful CRISPR/Cas9 double- strand breaks.

### Statistics and visualizations

GraphPad Prism 10 and R was used for graph visualizations and statistics. Unpaired t-tests were performed to determine significance between two groups. Correlations were calculated using Pearson’s Correlation. Three-way ANOVA analysis was used for navitoclax sensitivity between two groups at different doses. Gene signatures scores were received by calculating z-scores for each individual gene in a signature, followed by mean calculations, using Excel.

## List of abbreviations

BMP4:: bone morphogenetic protein 4
Cas9:: CRISPR-associated protein 9
CDK:: cyclin-dependent kinase
CDKN1A/2A:: cyclin-dependent kinase inhibitor 1A/2A
CL:: classical
CRISPR:: clustered regularly interspaced short palindromic repeats
EGF:: epidermal growth factor
FGF:: fibroblast growth factor
G1:: gap1 (cell cycle phase)
G2:: gap 2 (cell cycle phase)
GBM:: glioblastoma
KO:: knockout
MAPK:: mitogen-activated protein kinase
MES:: mesenchymal
OLIG2:: oligodendrocyte transcription factor 2
PI3K:: phosphatidylinositol 3 kinase
PN:: proneural
RB:: retinoblastoma transcriptional corepressor
SA-b-gal:: senescence-associated beta-galactosidase
SASP:: senescence-associated secretory profile
SMAD:: suppressor of mothers against decapentaplegic
SOX2:: SRY-box transcription factor 2
TGF-b:: transforming growth factor beta

## Declarations

### Ethics approval and consent to participate

Tumor sample collection of U3065MG cells was approved by the Uppsala regional ethical review board, number 2007/353 and informed consent was obtained from the patient. Experiments conformed to the principles set out in the WMA Declaration of Helsinki and the Department of Health and Humans Services Belmont Report.

### Consent for publication

Not applicable.

### Data availability

Bulk RNA sequencing data will be available upon request.

## Authors contributions

MN, ED, and BW designed the study. MN conducted all experiments and analyzed data. MN and ED composed the figures. ED and VR performed bioinformatic work. MN and AS designed and analyzed dose-response experiments. MN, ED, and BW wrote the manuscript, and all authors edited the manuscript.

## Acknowledgements

We thank the Swedish Cancer Society and the Swedish Research Council for financial support. We also thank Bo Segerman for initial processing of RNA sequencing data, and the staff at the Clinical Chemistry and Pharmacology division, Uppsala University Hospital, for help with drug response experiments (Kristin Blom, Jakob Rudfeldt and Anders Åkerström) and Incucyte analysis support (Malin Berglund). Some flow cytometry analyses were performed at the BioVis Platform, SciLife Laboratories, Uppsala.

## Supplementary Table

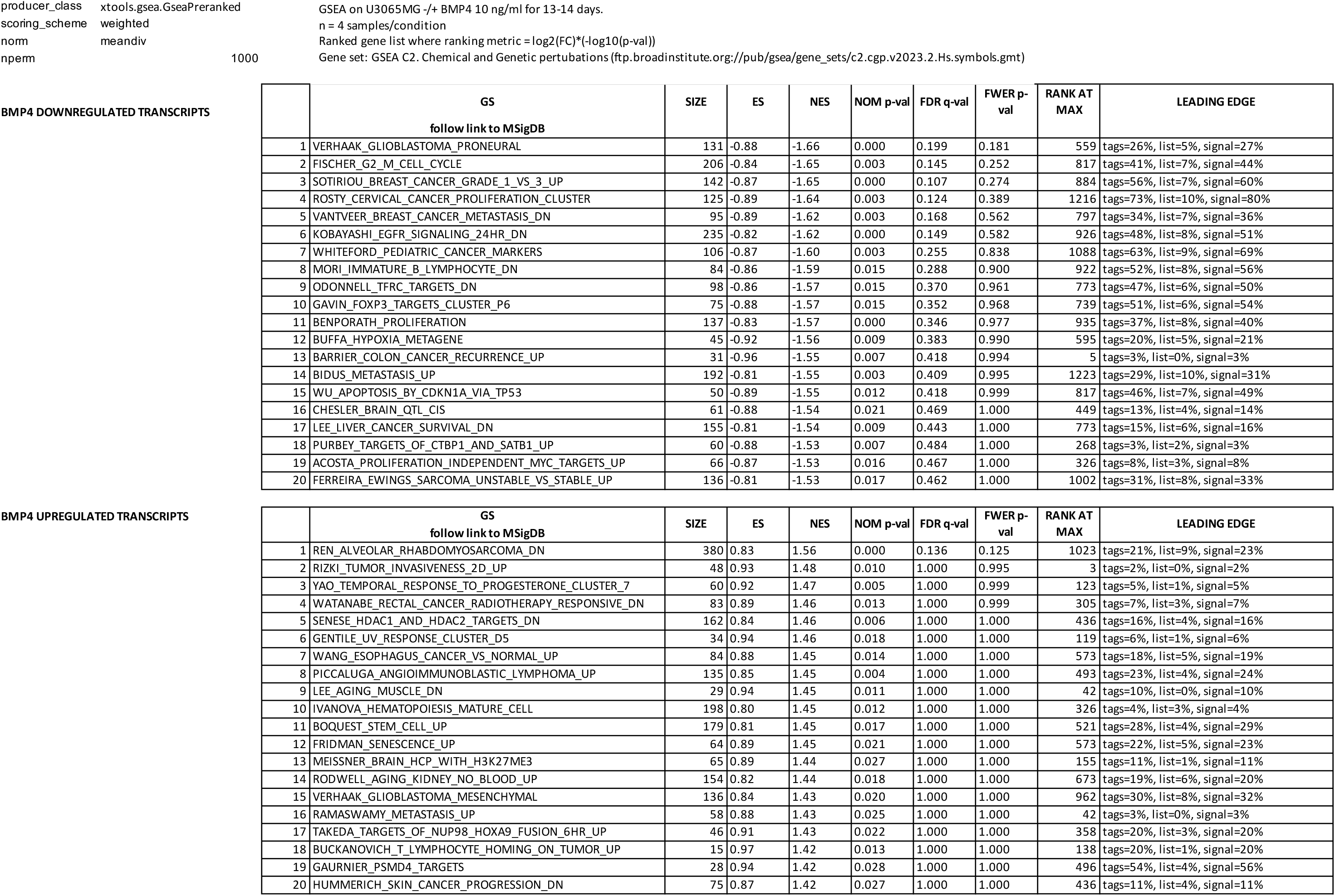

## Supplementary figures

**Supplementary Figure 1.**
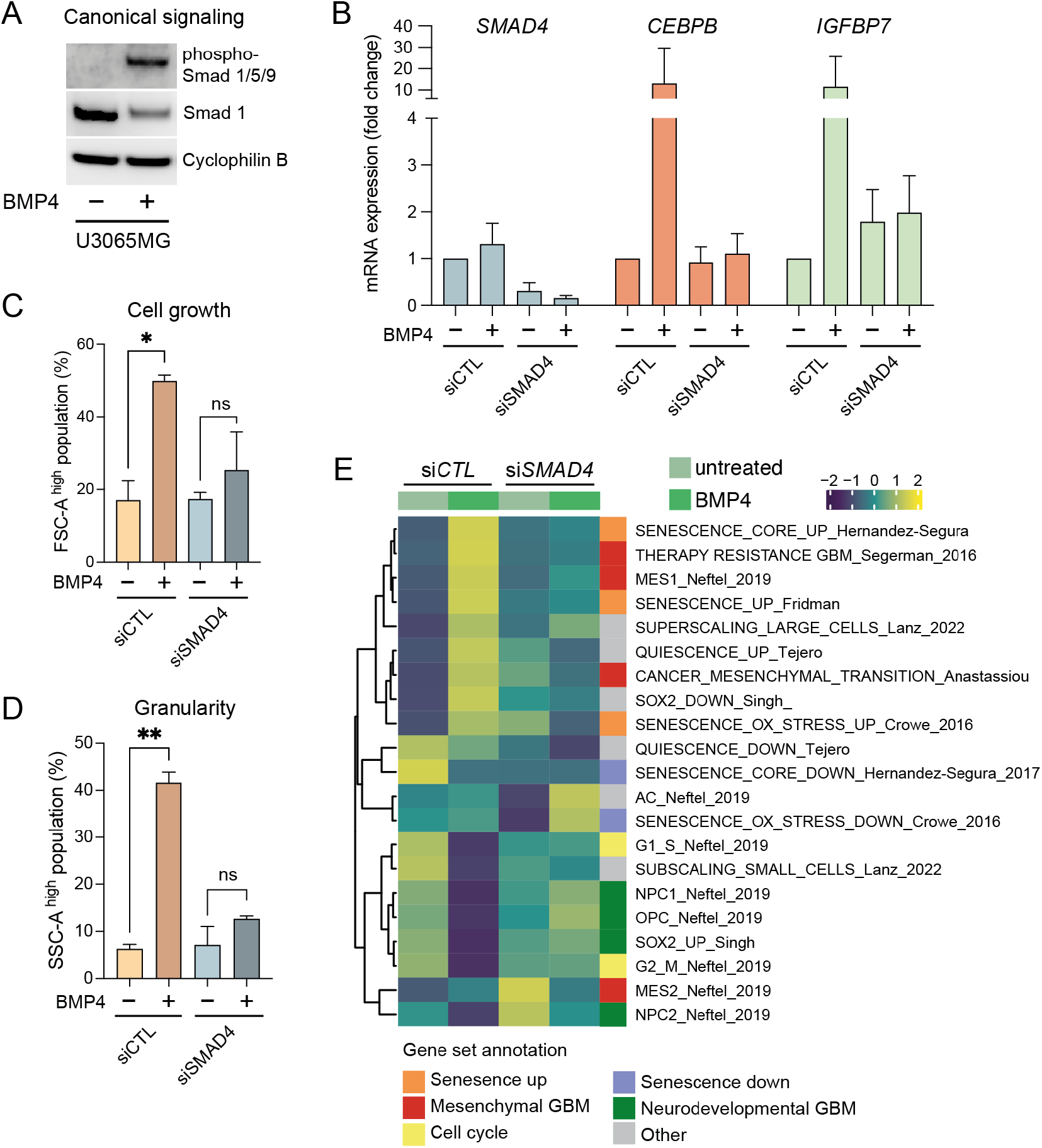
A. Phospho-SMAD1/5/9 and total SMAD1 western blot analysis of U3065MG cells stimulated with BMP4 (10 ng/ml) for 1 hour. B-E. siRNA-mediated knock-down of SMAD4 in U3065MG cells. Cells were transduced two times with siRNA against SMAD4 or scrambled control (siCTL) (day 0 and day 8), with addition of +/- BMP4 on day 2. BMP4 treatment for 12 days. B. mRNA expression of SMAD4, the senescence master-regulator gene CEBPB, and the SASP-related IGFBP7 using qRT-PCR analysis. C-D. Flow cytometry analysis of C) cellular size, gating on the forward-scatter-high cell population (FSC-A high), and D) cellular granularity, gating on the side scatter-high cell population (SSC-A high), as in Figure 1B-C. Two experiments, unpaired t-test, *, p=0.014; **, p=0.0024. E. Single-sample Gene Set Enrichment Analysis (ssGSEA) of transcriptome data from SMAD4 knockdown experiment.

**Supplementary Figure 2.**
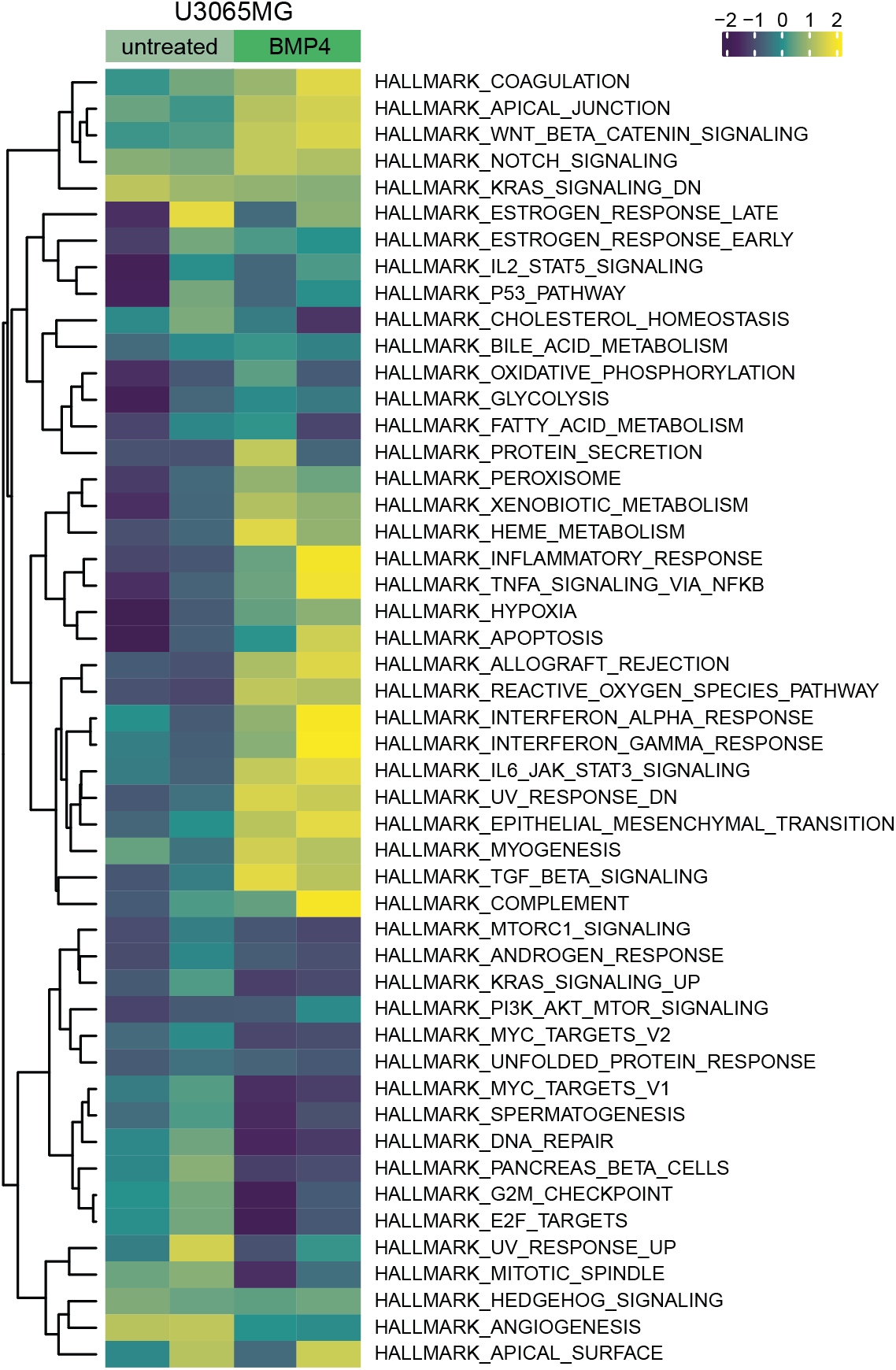
Heatmap of ssGSEA MSigDB Hallmark analysis on transcriptome data from untreated or BMP4-treated (14 days) U3065MG cells, two experiments.

**Supplementary Figure 3.**
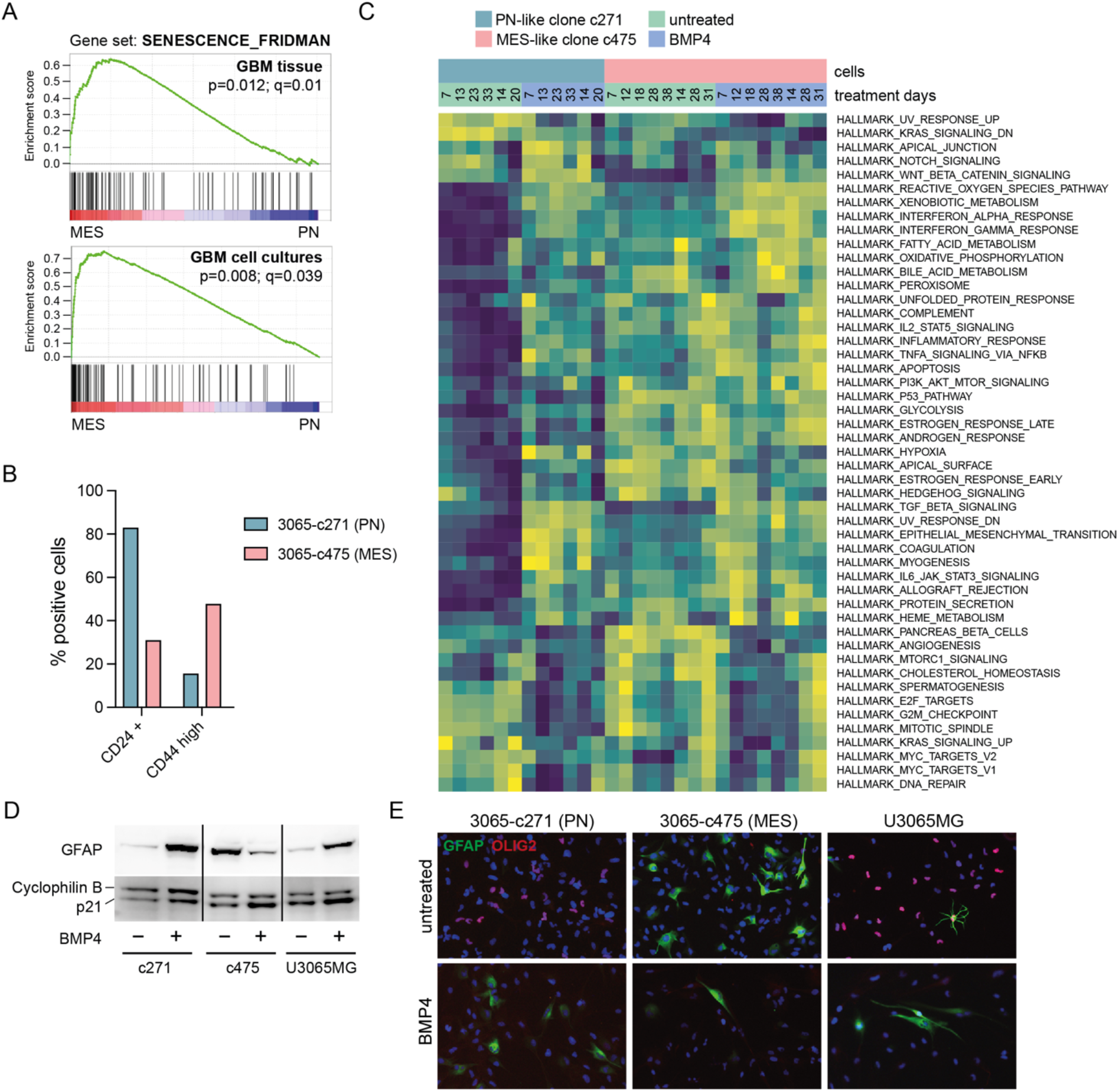
A. GSEA using the Fridman Senescence Up gene set on mesenchymal (n=158) vs. proneural (n=139) GBM tumors from TCGA database (top panel); and 19 mesenchymal vs. 14 proneural GBM cell lines from the HGCC repository (bottom panel). B. Bar graph showing percentage of CD24 and CD44-high expressing cells in PN-like 3065-c271 and MES-like 3065-c475 cells using flow cytometry analysis. C. Single sample GSEA using MSigDB Hallmarks gene sets in untreated and BMP4-treated PN-like clone 3065-c271 and MES-like clone 3065-c475. D-E. GFAP western blot (D) and OLIG2 and GFAP immunofluorescence staining (E) of U3065MG, 3065-c271 and 3065-c475 treated +/-BMP4 for two weeks.

**Supplementary Figure 4.**
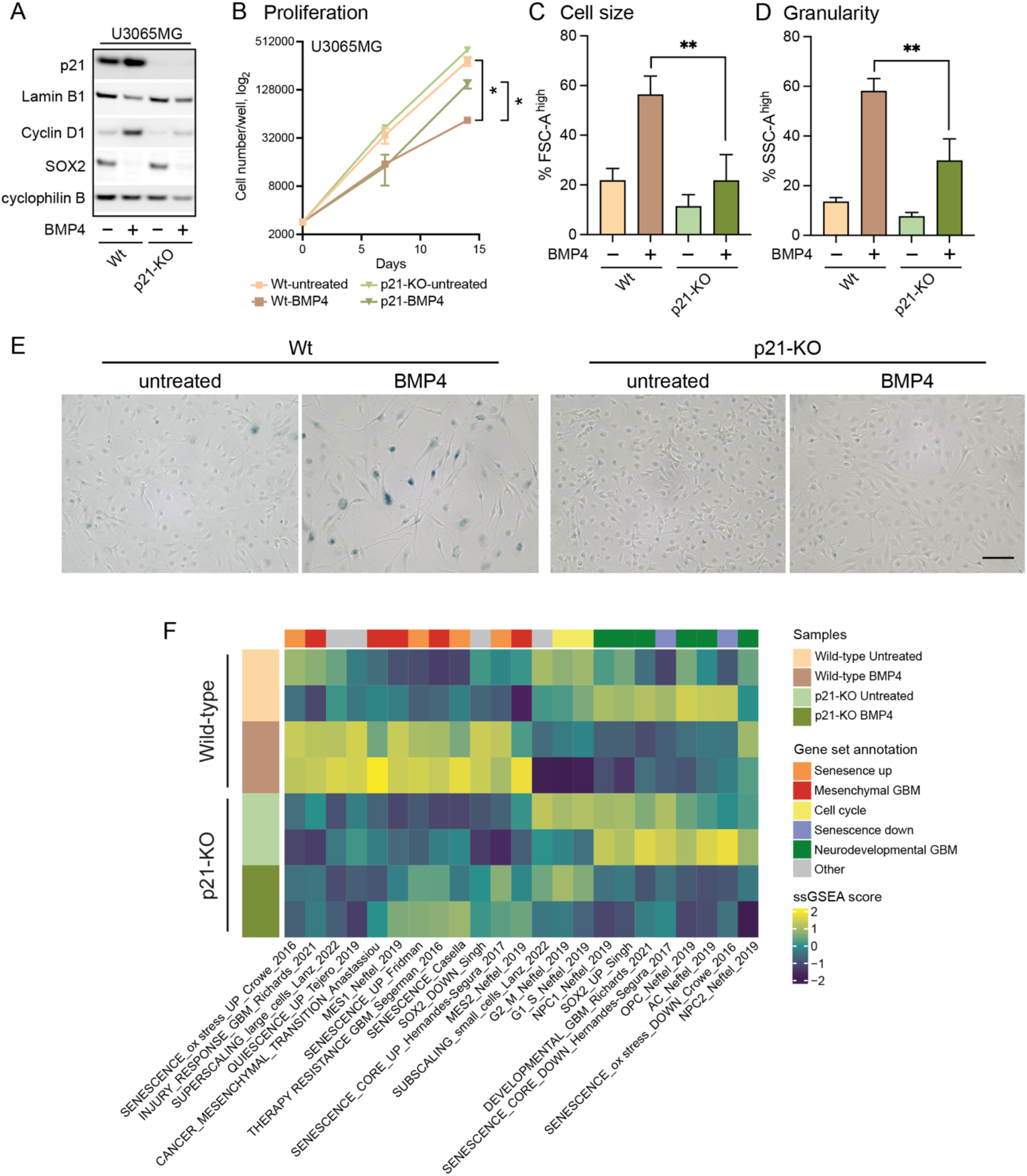
Western blot on wild-type and p21 knockout (p21-KO) U3065MG cells +/- BMP4 using antibodies against p21, lamin B1, cyclin D1, SOX2, and cyclophilin B. B. Proliferation of wild-type and p21-KO cells +/- BMP4. Cell counting on day 0, day 7 and day 14. C. Flow cytometry analysis of cell size (cell growth) using forward scatter (FSC-A) measurements and quantification of the FSC-A high cell population; and cell granularity (D). Side scatter area (SSC-A) high gated population is plotted. E. SA-β-gal staining, scale bar 100 µm. F. Heatmap of ssGSEA-scores of U3065MG wild-type and p21-KO cells +/-BMP4.

**Supplementary Figure 5.**
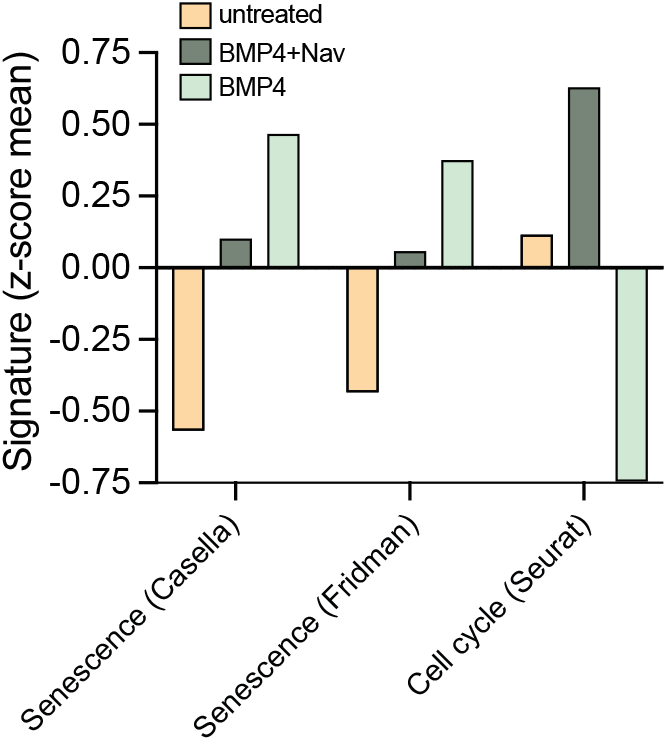
Bar graph showing senescence (42, 44) and cell cycle (57) gene signature scores (mean z values) of untreated, and BMP4-treated +/- navitoclax 3065-c475 cells.

